# Sleep ripples drive single-neuron reactivation for human memory consolidation

**DOI:** 10.64898/2026.03.27.714528

**Authors:** Marcel S. Kehl, Thomas P. Reber, Valeri Borger, Rainer Surges, Florian Mormann, Bernhard P. Staresina

**Affiliations:** Department of Epileptology, University Hospital Bonn, Bonn, Germany; Department of Experimental Psychology, University of Oxford, Oxford, UK; Oxford University Centre for Integrative Neuroimaging, University of Oxford, Oxford, UK; Faculty of Psychology, UniDistance Suisse, Brig, Switzerland; Department of Neurosurgery, University Hospital Bonn, Bonn, Germany

**Keywords:** Hippocampus, SWR, Ripple, Human Single Neurons, Memory, Consolidation, Sleep, Medial Temporal Lobe

## Abstract

Sleep transforms fragile experiences into lasting memories, but the neuronal basis of this process in humans has remained elusive. In rodents, hippocampal ripples orchestrate the replay of place cell sequences, establishing a cellular mechanism for consolidation – though with limited generalizability to human memory. In humans, neuroimaging has revealed large-scale offline reactivation, but these coarse signals leave open whether individual neurons are reactivated and how ripples might mediate this process. Here, we bridge this gap by directly recording 1,466 medial temporal lobe (MTL) neurons and intracranial electroencephalography during learning, post-learning wakefulness, and sleep. We show that ripples robustly drive neuronal firing, with sleep ripples eliciting stronger activation than wake ripples. Critically, neurons tuned to items that were later remembered fired more strongly during ripples than those coding for forgotten items, and this memory-linked reactivation was selectively observed during sleep. Finally, ripple-associated neuronal MTL bursts were detectable across widespread cortical activity, pointing to a mechanism for systems-level consolidation. Together, these findings provide the first direct evidence that ripple-driven single-neuron reactivation supports human episodic memory consolidation and reveal why sleep — compared to wakefulness — offers a privileged window for stabilizing memories.

## Main Text

The ability to stabilize fragile new experiences into enduring memories is fundamental for learning and cognition. Sleep is particularly effective in supporting this process (*1–3*), yet the cellular mechanisms by which sleep promotes memory consolidation in humans are not known.

In rodents, hippocampal sharp-wave ripples — brief bursts of high-frequency activity during non-REM sleep (*4*) — coordinate the replay of place cell sequences, establishing a cellular mechanism for memory consolidation (*5–7*). Selective disruption of ripple events impairs subsequent recall (*8*), underscoring their causal role. Yet it remains unclear whether and how such replay of spatial navigation patterns relates to the intricate phenomenon of human episodic memory (*9*).

In humans, neuroimaging has revealed large-scale reactivation of memory traces during post-learning rest and sleep (*10–14*), and intracranial recordings have identified ripple events in the MTL and beyond (*15–19*). These observations point to ripples as a candidate mechanism for human memory consolidation, but they leave open two key questions. First, there is no evidence that individual human neurons holding mnemonic content are reactivated during ripples, leaving a gap between rodent replay models and human systems-level findings. Second, ripples occur during both wakefulness and sleep (*20–23*), yet sleep promotes long-term retention more effectively. Whether ripple-triggered neuronal reactivation differs between these states, and whether such differences explain the privileged role of sleep, has not been established.

Here, we bridge these gaps by directly recording ripples as well as firing rates from 1,466 MTL neurons across learning, post-learning wakefulness, and sleep. We show that ripples robustly drive neuronal firing, that sleep ripples preferentially recruit neurons coding for subsequently remembered items, and that ripple-associated bursts reverberate across distributed cortical networks. Together, these findings provide direct single-neuron evidence that ripple-driven reactivation underlies human memory consolidation and reveal why sleep offers a privileged window for stabilizing memory.

## Results

### Tracking neurons across learning and sleep

We capitalized on the rare opportunity to record single-neuron activity in patients undergoing invasive monitoring for drug-resistant epilepsy (Fig. 1A). Across 14 overnight sessions in 9 participants, we simultaneously obtained polysomnography, intracranial EEG, and microwire recordings from MTL neurons (Fig. 1B). This approach yielded 1,466 neurons, whose spiking activity could be aligned with local ripple events. An example trace shows how a hippocampal ripple — marked by a brief high-frequency oscillation followed by a sharp wave — coincides with a burst of neuronal firing (Fig. 1A).

**Fig. 1.**
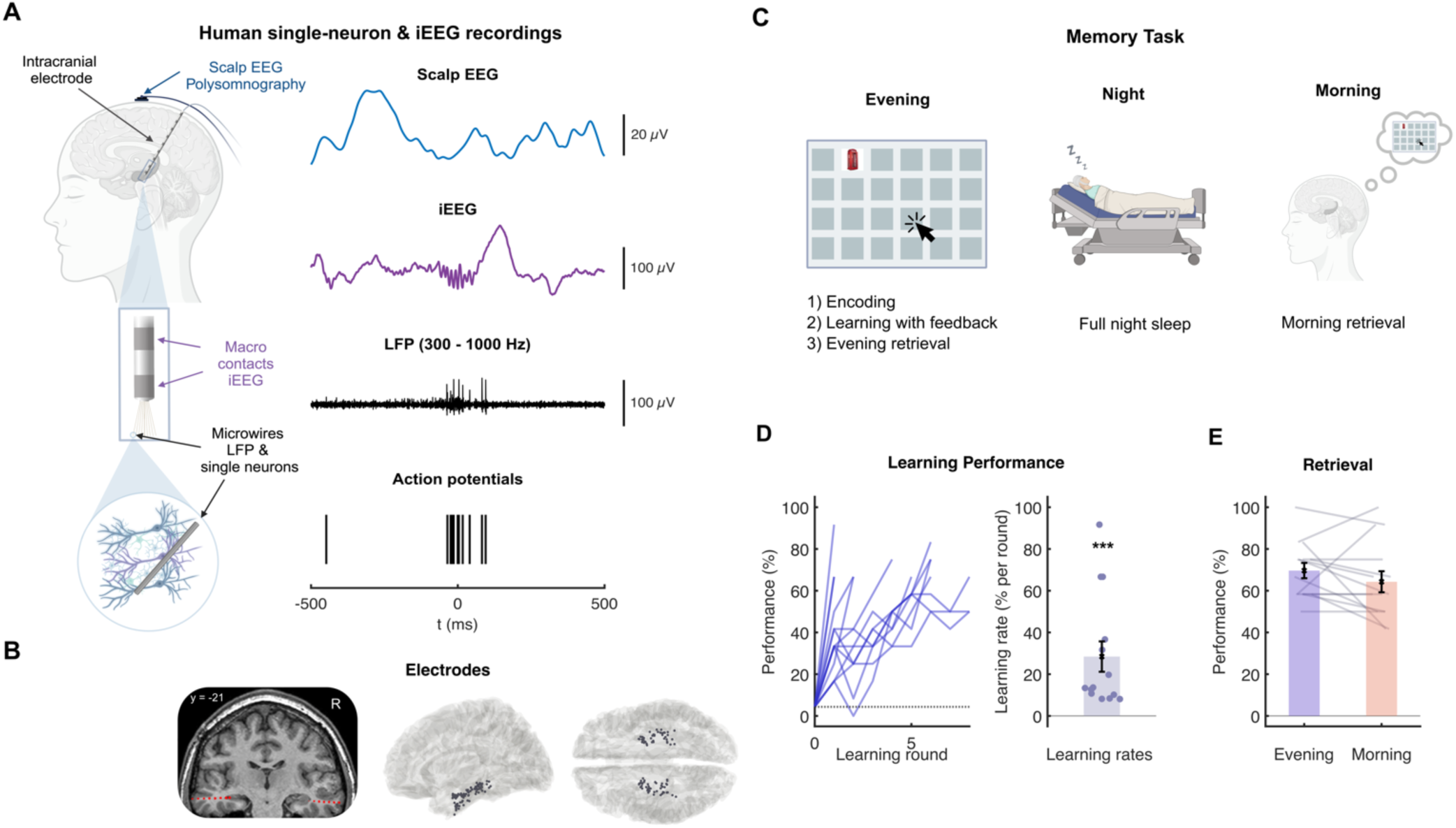
Tracking neurons across learning and sleep. **(A)** Simultaneous scalp EEG, intracranial EEG, local field potentials (LFP), and single-neuron recordings in patients undergoing presurgical epilepsy monitoring. Scalp EEG was used for sleep staging (blue, top). Ripple events were detected from MTL macro contacts (purple, second row). Microwire bundles recorded LFPs and action potentials (black, third and fourth row). Created in BioRender. Kehl, M. S. (2026) https://BioRender.com/5v44xhw **(B)** Left: Co-registered CT/MRI scans for visualizing electrode locations; Right: Black dots mark innermost electrode positions projected to MNI space across participants. **(C)** Memory task. Before sleep, participants learned the positions of 12 image pairs on a grid. Each pair was presented twice during encoding, followed by repeated study–test rounds with feedback. Learning ended once ≥8 pairs (66%) were recalled or after 8 blocks. Retrieval was tested without feedback in the evening, followed by overnight sleep and a morning retrieval task. Created in BioRender. Kehl, M. S. (2026) https://BioRender.com/gpklzsv **(D)** Learning performance (blue lines, one per recording session) increased across blocks, significantly above chance (28 ± 7.3% per round, *P* = 0.00012, *N* = 14, two-sided Wilcoxon signed-rank test vs. chance). Chance was defined as 1 in 23 positions (4.35%). **(E)** Retrieval performance (mean ± s.e.m.) in the evening (70 ± 3.7%) and the following morning (64 ± 5.1%; *P* = 0.26, *N* = 14, two-sided Wilcoxon signed-rank tests versus chance).

Participants engaged in a learning task in which they memorized the locations of 12 image pairs (Fig. 1C). After repeated study–test rounds with feedback, they completed an evening retrieval test without feedback, followed by a night of sleep with polysomnography. Memory was probed again the next morning, showing robust overnight retention (Fig. 1D–E). This design enabled us to track MTL neurons from encoding through post-learning wake and sleep.

### Ripple-locked firing of human MTL neurons

After identifying and characterizing ripple attributes during sleep and awake periods (fig. S1), we examined how spiking of individual MTL neurons aligns with ripples. Fig. 2A shows an example hippocampal neuron that markedly increased its firing around ripples during sleep compared to wake, illustrating a clear state-dependent effect. To test this at the population level, we computed ripple-locked firing rates for all 1,466 recorded neurons. On average, sleep ripples elicited a sharp increase in firing followed by a brief suppression, whereas wake ripples produced a weaker increase without subsequent suppression (Fig. 2B). Direct within-neuron comparisons confirmed that firing rates were significantly higher during sleep ripples than during wake ripples (Fig. 2C). Thus, although ripples in both states engage MTL neurons, sleep ripples are associated with stronger and more temporally structured activation.

**Fig. 2.**
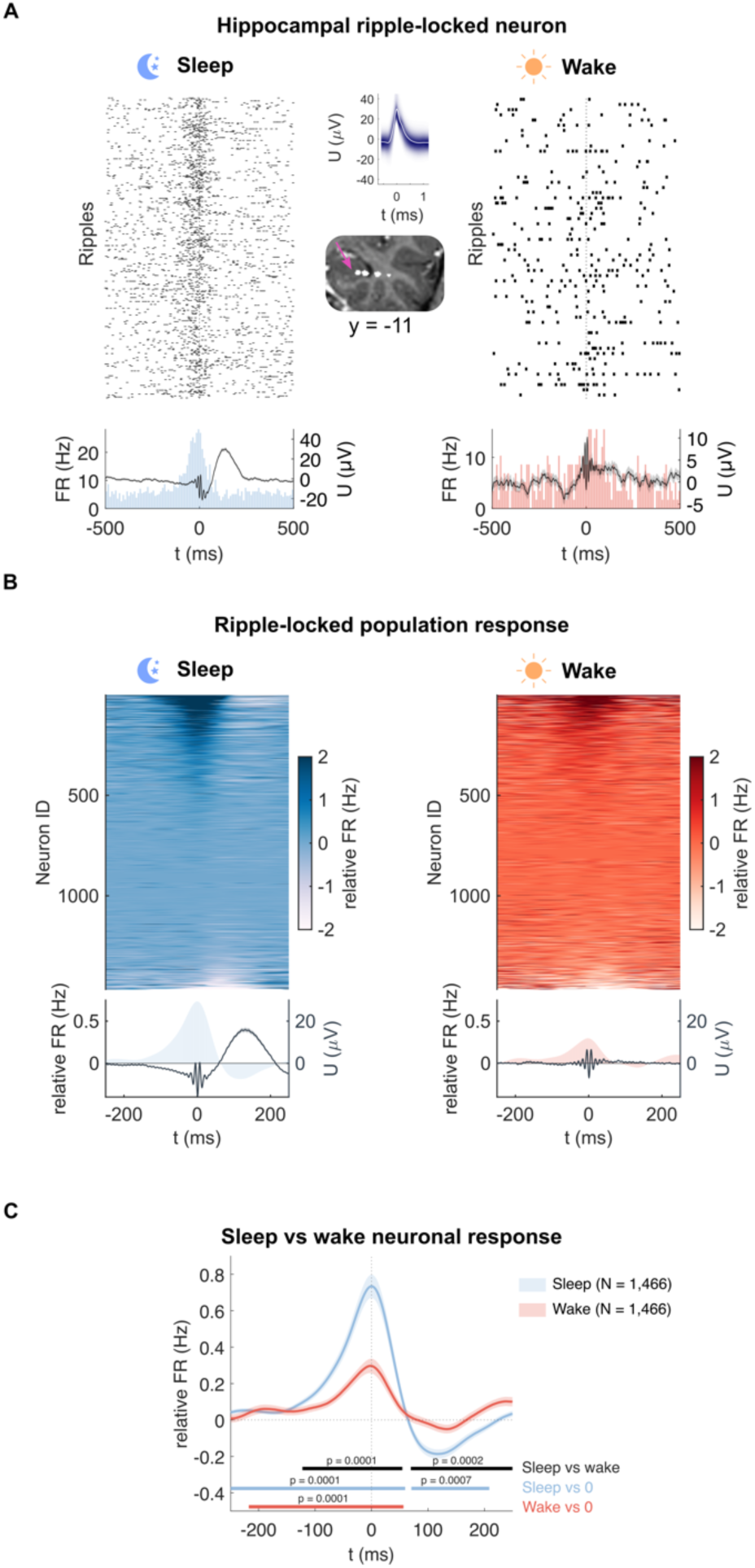
Ripple-locked firing of human MTL neurons. **(A)** Raster plot and peristimulus time histogram (PSTH) of a hippocampal neuron aligned to ripple onset during sleep (left) and wake (right). The neuron showed a stronger increase in firing around sleep ripples than wake ripples. **(B)** Population activity aligned to ripple onset during sleep (left) and wake (right). Each row represents one neuron’s firing-rate (FR) change (normalized to baseline, smoothed with a 100-ms Gaussian kernel, neurons sorted by mean FR). The mean across all 1,466 neurons is shown below, together with the average ripple waveform. **(C)** Mean population firing rates (± s.e.m.) during sleep (blue) and wake (red) ripples. Both states showed significant increases relative to baseline (horizontal bars), but firing was significantly stronger during sleep ripples (black bar; cluster-permutation tests, 10,000 permutations).

### Sleep ripples preferentially recruit neurons coding for remembered stimuli

The human MTL contains neurons that fire selectively to specific concepts, such as an object or a person (“concept cells”; (*24–27*)). These sparse, stimulus-specific, semantically invariant, and context-independent representations (*28*) have been hypothesized to form the semantic building blocks of episodic memory (*29*), a hypothesis only recently confirmed experimentally (*30*). Here, these stimulus-selective neurons provide the unique opportunity to test whether ripple-locked reactivation targets the very neurons involved in encoding remembered experiences.

To identify stimulus-selective neurons, participants first viewed a large image set in a screening session, and learning stimuli for the current study were selected from images that evoked selective neuronal responses. Across all sessions, we recorded 91 selective neurons, each responding to exactly one of the 12 learning images. These neurons allowed us to link neuronal reactivation during ripples directly to subsequent memory outcomes (Fig. 3A).

**Fig. 3.**
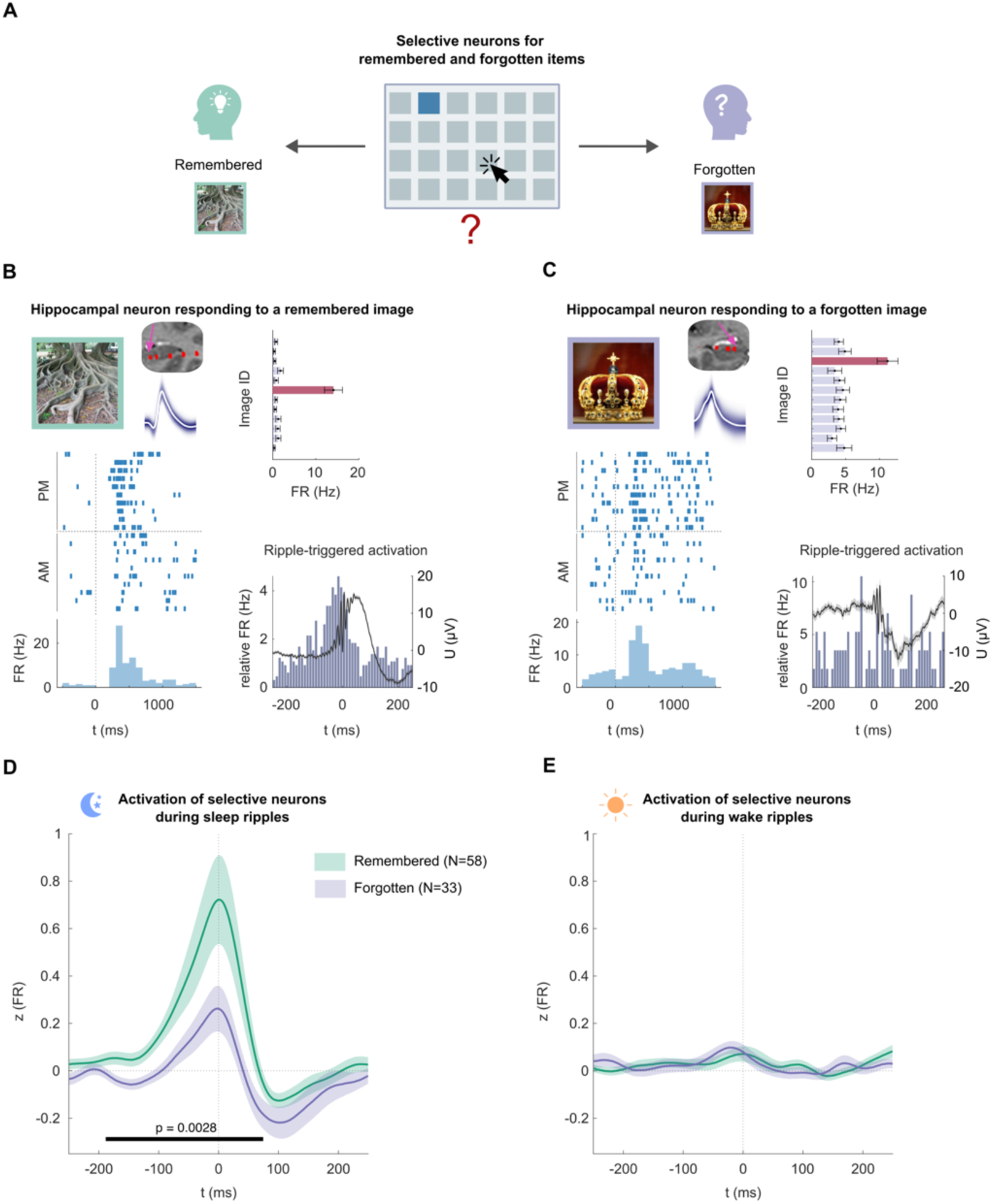
Sleep ripples preferentially recruit neurons coding for remembered stimuli. **(A)** Overall, 91 neurons responded selectively to one of the 12 learning images, enabling a direct link between content-specific ripple-locked activity and memory outcome. Stimulus-selective neurons were categorized based on whether images were correctly remembered across both retrievals (evening and morning) or not (N_Remembered_ = 58, N_Forgotten_ = 33). **(B)** Example hippocampal neuron selective for the image of tree roots (later remembered). Left: Raster Plot and PSTH showing selective responses to the preferred image during screening sessions conducted the evening before and the morning after the main experiment. Right top: Firing rates (mean ± s.e.m., 0 to 500 ms) to all 12 images, with the preferred stimulus marked in red. Right bottom: The same neuron showed a sharp increase in firing around sleep ripples. **(C)** Example hippocampal neuron selective for the image of a crown (later forgotten). This neuron responded selectively during screening but did not increase firing around sleep ripples. **(D)** Population ripple-locked activity of neurons selective for remembered (green, *N* = 58) versus forgotten (blue, *N* = 33) items during sleep. Neurons coding for remembered stimuli showed significantly stronger activation around ripples (*P* = 0.0028, cluster-permutation test, 10,000 permutations). **(E)** During wake ripples, no significant difference was observed between neurons coding for remembered vs. forgotten stimuli.

Single-neuron examples reveal distinct activation patterns during ripples. For instance, a hippocampal neuron selective for the image of tree roots (later successfully recalled) showed a sharp increase in firing around sleep ripples (Fig. 3B). By contrast, a neuron selective for the image of a crown (later forgotten) showed no such ripple-linked increase (Fig. 3C). At the population level, neurons selective for subsequently remembered stimuli were significantly more active during sleep ripples than neurons selective for forgotten stimuli (Fig. 3D). Crucially, this difference was observed only during sleep ripples, not wake ripples (Fig. 3E). Direct comparison of sleep versus wake ripple responses confirmed that memory-selective reactivation was specific to sleep (fig. S2A). Finally, no such activation difference was observed during ripple-free, surrogate sleep events (fig. S2B).

Together, these results demonstrate that sleep ripples recruit neurons coding for learned content and that the degree of such recruitment predicts successful item-specific memory, revealing a cellular mechanism by which sleep supports human memory consolidation.

### Ripple-linked bursts couple MTL neurons to cortical networks

The tight link between ripple events and increased firing of neurons coding for remembered stimuli creates temporally structured activity patterns well suited for synaptic plasticity (*31*, *32*). However, long-term stabilization is thought to also depend on *systems consolidation*—the transfer of mnemonic information from hippocampus to cortex (*2*, *33*, *34*). We therefore asked whether ripple-triggered neuronal activity is also broadcast beyond the MTL (Fig. 4A).

**Fig. 4.**
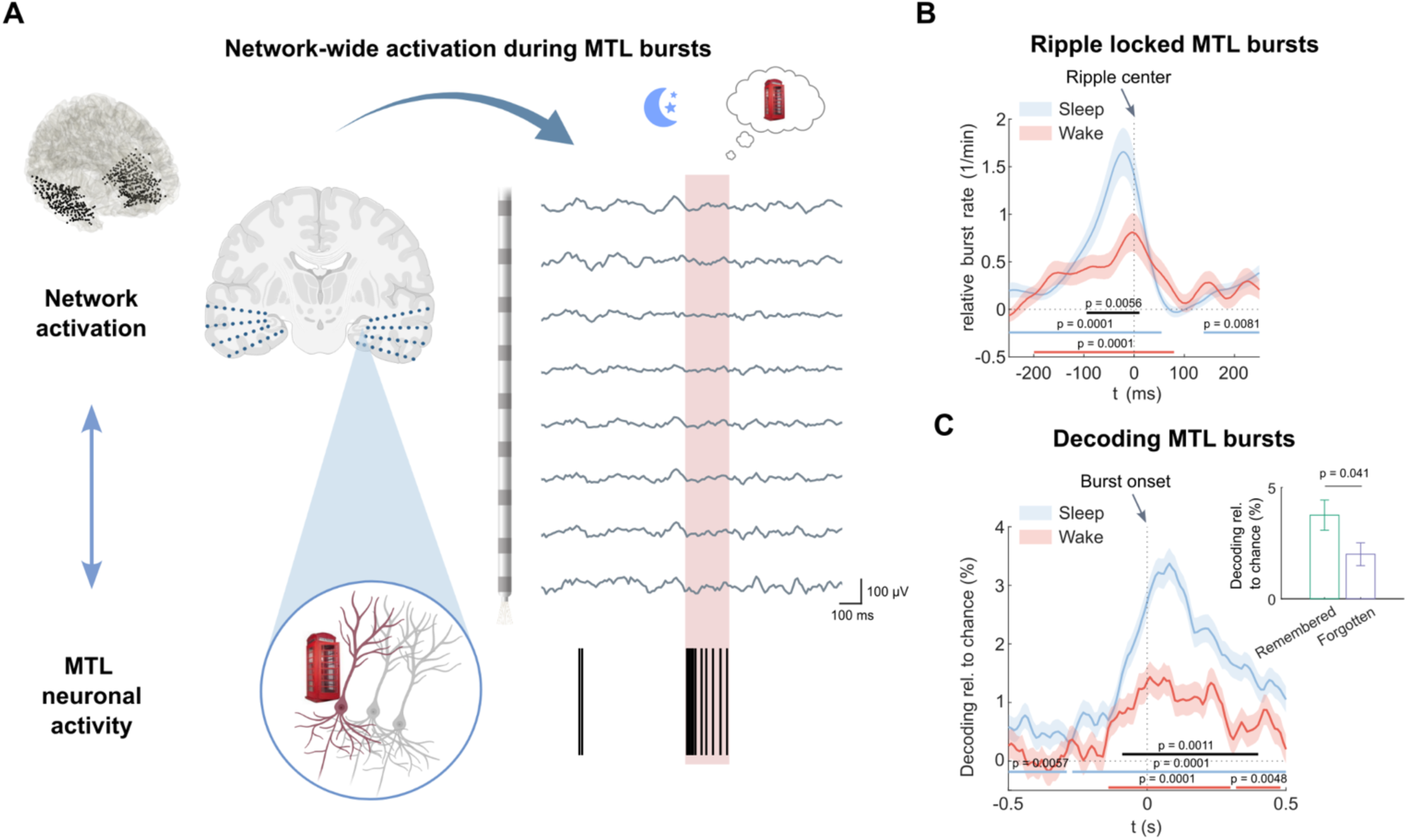
Ripple-linked bursts couple MTL neurons to cortical networks. **(A)** Top left: Overview of macro-contacts (black dots) across patients, projected onto MNI space, covering ventral and lateral temporal cortex. Simultaneous single-neuron and intracranial EEG recordings enable linking local MTL firing to widespread cortical activity. Right: Illustration of a depth electrode with eight macro contacts recording iEEG activity synchronized with the spiking of a stimulus-selective neuron (“phone booth”). The red-shaded interval marks a burst of neuronal activity. Created in BioRender. Kehl, M. S. (2026) https://BioRender.com/ufznb43 **(B)** Burst rates (mean ± s.e.m.) increased significantly around ripples compared to baseline during both wakefulness and sleep (blue and red horizontal bars, cluster-permutation test, 10,000 permutations, N = 1,466 neurons), demonstrating robust ripple–burst coupling. Sleep ripples were associated with stronger burst responses than wake ripples (black horizontal bar). **(C)** Decoding of burst events from distributed cortical iEEG activity. Classifiers distinguished real bursts from burst-free controls during both wake (red) and sleep (blue). Decoding was significantly above chance in both states (blue and red horizontal bars), but significantly higher during sleep than wake (black bar, cluster-permutation test, 10,000 permutations, N = 791 neurons with sufficient burst events). Burst decoding was significantly higher for neurons selective for remembered compared to forgotten items (inset bar graph, mean ± s.e.m., Wilcoxon rank-sum test of mean decoding accuracy during significant time interval (black), *P* = 0.041, N = 88 neurons with sufficient burst events: 56 remembered vs. 32 forgotten).

As a first step, we focused on neuronal bursts, brief episodes of elevated spiking that are potent drivers of synaptic plasticity (*35*, *36*). Using a conservative burst-detection algorithm (fig. S3) (*37*), we found that burst rates increased sharply around ripples (Fig. 4B). This increase was again more pronounced during sleep, indicating that sleep ripples provide privileged windows for burst generation.

To assess whether these local bursts propagate to wider brain networks, we leveraged simultaneous single-neuron and intracranial EEG recordings across distributed cortical sites (Fig. 4A). Multivariate classifiers trained on cortical iEEG could reliably distinguish time windows containing MTL bursts from matched, burst-free control periods (Fig. 4C). In other words, discrete single-neuron bursts are associated with detectable signatures at the macroscopic network level. Critically, decoding accuracy was significantly higher during sleep than during wakefulness, indicating stronger coupling between local MTL firing and large-scale cortical activity during sleep (Fig. 4C). Moreover, bursts from neurons selective for remembered items could be decoded significantly better at the network level than those from neurons selective for forgotten items (Fig. 4C). These findings suggest that ripple-triggered bursts not only strengthen local memory traces but also promote the transfer of information to neocortex, providing a potential cellular-to-systems pathway for memory consolidation.

## Discussion

Our findings provide the first direct evidence that reactivation of neurons representing the content of recent experiences during sleep ripples in the human medial temporal lobe predicts successful memory formation. By linking ripple activity to both single-neuron selectivity and subsequent memory outcomes, we identify a cellular mechanism for human memory consolidation and explain why sleep provides a privileged window for this process. In rodents, hippocampal ripples coordinate the replay of place cell sequences, but whether and how this phenomenon relates to human episodic memory remains elusive (*38*). In humans, neuroimaging has revealed offline reactivation of large-scale memory representations (*12*, *39*, *40*), but the underlying neuronal mechanisms remain unknown. By showing that the recruitment of stimulus-selective neurons during ripples in the human MTL predicts subsequent memory of their preferred stimuli, our results bridge these two strands of evidence.

A key advance is that this memory-linked reactivation was specific to sleep ripples. While ripples occur both during wakefulness and sleep (*15*, *19*, *22*, *41*), only sleep ripples robustly boosted neuronal firing of neurons selective to remembered items. This sleep specificity provides a mechanistic explanation for the long-standing observation that sleep supports memory consolidation more effectively than wake (*2*, *42*). Ripple-triggered activation generates temporally precise firing patterns conducive to synaptic plasticity (*36*, *18*, *43*, *44*), positioning sleep ripples as privileged windows for strengthening memory traces.

At the same time, memory consolidation is thought to involve not only local strengthening but also the transfer of memory representations to neocortex (*45*, *46*, *1*, *2*). Our data show that ripple-triggered bursts of MTL neurons could be reliably decoded from widespread cortical activity, and this coupling was again stronger during sleep than wake and stronger for neurons coding for remembered vs. forgotten items. These findings suggest that ripples do not merely reactivate local memory-related neurons, but also broadcast their activity across distributed cortical networks, providing a plausible mechanism for systems consolidation in the human brain.

Several questions remain. Our paradigm did not test for sequential replay, a hallmark of rodent studies, and future work with navigation or sequence learning tasks will be critical to determine whether human ripple-linked reactivation reinstates learned temporal patterns (*38*). Likewise, our recordings sampled only a small fraction of MTL neurons and cortical regions, and higher-density approaches will be needed to capture the full population dynamics. Finally, causal manipulations of ripple events — for example, using closed-loop stimulation to enhance or disrupt reactivation (*8*, *20*, *47*) — will be essential to establish the necessity of ripple-triggered activity for human memory consolidation.

Together, our findings uncover how ripple-triggered reactivation at the level of single neurons contributes to human memory consolidation. By showing that sleep ripples preferentially recruit neurons coding for remembered experiences and broadcast their activity to cortex, our study links rodent replay and human neuroimaging findings within a unified cellular framework. Sleep emerges not simply as a passive state for memory retention, but as an active window in which ripples stabilize fragile traces and embed them into distributed cortical networks, providing a mechanistic account of sleep-dependent memory consolidation.

## Acknowledgments

We thank the patients for their participation. Icons used in figures were obtained through an Icon Pro license from the Noun Project (https://thenounproject.com).

## Funding

European Research Council (ERC) via the European Union’s Horizon 2020 grant 101001121 (BPS)

German Research Foundation (DFG, MO 930/4-2, MO 930/15-1, SPP 2411) (FM)

Ministry of Culture and Science of the State of North Rhine-Westphalia (iBehave Network) (FM)

## Author contributions

MSK, FM, BPS and TPR designed the study. RS and FM recruited patients. VB and FM performed the surgeries and implanted the electrodes. MSK conducted all experiments and pre-processed the data. MSK, BPS and FM planned the analyses. MSK analysed the data with input from BPS. MSK and BPS designed and prepared the figures. FM and BPS supervised and administered the project and secured funding. MSK, BPS and FM wrote the manuscript. All authors discussed the results and commented on the manuscript.

## Data and materials availability

Data and analysis code used to generate the main figures and to support the findings of this manuscript will be made publicly available upon publication.

## List of Supplementary Materials

Materials and Methods

Figs. S1 to S3

Table S1

## Supplementary Materials

### Materials and methods

#### Participants

We analyzed 14 recording sessions from 9 patients with drug-resistant epilepsy undergoing invasive seizure monitoring at the Department of Epileptology, University of Bonn Medical Center (42.6 ± 12.6 years, mean ± s.d.; 5 female, 4 male; table S1). All patients were evaluated for potential neurosurgical treatment and implanted with depth electrodes for clinical purposes. All participants gave written informed consent, and the study was approved by the University of Bonn Medical Institutional Review Board.

#### Single-neuron recordings

Electrophysiological signals were acquired with a 256-channel ATLAS amplifier and Pegasus software (v2.1.1, Neuralynx, Bozeman, MT). All patients were implanted with Behnke–Fried depth electrodes (AdTech, Racine, WI) containing eight cylindrical macro-contacts (innermost spacing 3 mm; Fig. 1A). Each electrode housed a bundle of eight high-impedance platinum–iridium microwires and one non-insulated reference wire, extending 3–5 mm beyond the electrode tip. Implantation target locations were determined solely by clinical needs, and correct electrode placement was ensured by stereotactic surgery and postoperative CT. Electrodes were implanted bilaterally and targeted MTL structures including hippocampus, amygdala, entorhinal cortex, and parahippocampal cortex. Electrodes not targeting these MTL regions were not analyzed.

Signals were sampled at 32,768 Hz, band-pass filtered (0.1–9,000 Hz), and spike-sorted across the entire recording using Combinato (*48*), a toolbox optimized for long-term human single-unit recordings. Sorting results were further curated by an experienced annotator to remove artifacts and merge clusters based on spike waveform shapes, inter-spike intervals, and cross-correlograms. Across all sessions, we identified 1,466 neurons in the MTL (of which 789 were classified as single units and 677 as multi-units).

#### Experimental paradigm

The memory paradigm began in the evening, 1–2 hours before participants’ habitual bedtime. Participants were asked to memorize the positions of 24 cards arranged in a 4 x 6 grid on a computer screen (*39*). For each session, images were selected individually based on their ability to elicit selective neuronal responses in a prior screening on the same day, during which participants viewed a large set of images (*24*, *49*, *50*).

During an initial learning phase, all card pairs were presented twice in two consecutive blocks of pseudorandom order: the first card appeared for 1 s, followed by its paired card. Participants were instructed to memorize the pair location and proceed to the next trial with a button press. In the subsequent learning-with-feedback phase, only the first card was shown, and participants had to select the corresponding paired card using the computer cursor. Choices were followed by visual feedback (green/red frame) and the correct card pair was displayed for 2 s. The 12 pairs were repeated in blocks of pseudo-random order until participants either recalled ≥8 card locations correctly or completed 8 blocks. After learning, participants completed a retrieval block without feedback. Next morning (after a full night of sleep), participants completed a second retrieval block without feedback. The morning retrieval occurred on average approximately 9.5 hours after the evening session (567.5 ± 26.4 min, mean ± s.e.m.). Our analysis was restricted to ripple events occurring after completion of the evening task and before morning task onset. During this interval, participants remained awake for approximately 3 hours (177.7 ± 32.2 min) and spent about 4.5 hours in non-REM stages N2 and N3 (269.2 ± 21.3 min).

The memory task was part of a contextual cueing paradigm, where each card was paired with one of three odors during encoding. During N2 and N3 sleep, two encoding odors and a novel control odor were presented intermittently in 30-s blocks. For the present study, all sleep periods containing odor presentations were excluded from analysis, and no significant performance differences were observed between cued and uncued items linked to stimulus-selective neurons.

#### iEEG recordings and data preprocessing

All analyses were performed in MATLAB 2024b using the FieldTrip toolbox (*51*). Single-neuron, iEEG, and scalp EEG signals were amplified and recorded simultaneously on a 256-channel ATLAS system (Neuralynx, Bozeman, MT), ensuring precise temporal alignment across signals. iEEG data were downsampled to 1,000 Hz and locally re-referenced by subtracting the signal of the second innermost macro-contact from the innermost (bipolar referencing). Signals were high-pass filtered at 0.1 Hz and notch-filtered at 50-Hz harmonics (49–51, 99–101, 149–151, 199–201 Hz). Following established protocols (*18*), artifacts were automatically detected separately for each vigilance stage (wake, N1, N2, N3, REM). Data segments were marked as artifacts when amplitude, as well as the gradient, or high-frequency activity (>250 Hz) exceeded 5 median absolute deviations (MAD), or when any of these features alone exceeded 12 MAD. Interictal epileptiform discharges were identified with an automated algorithm (*52*) and excluded. All detected artifacts were padded with a ±1 s buffer, and intervals shorter than 3 s between consecutive artifacts were discarded.

#### Sleep scoring and ripple detection

Alongside the intracranial recordings, polysomnographic (PSG) data were collected using the same amplifier, including scalp Electroencephalography (EEG channels: C3, C4, F3, F4, O1, O2), electrooculography (EOG) and chin electromyography (EMG). PSG data were used for sleep scoring with YASA (*53*) and SomnoBot (https://somnobot.fh-aachen.de), the latter of which is an implementation of RobustSleepNet (*54*). An experienced annotator manually validated and curated all automated sleep scoring results with special emphasis on epochs in which the automated algorithms showed disagreement.

Ripples were detected using established detection methods (*18*, *55*). In brief, the iEEG signal was bandpass filtered between 80 and 120 Hz (using FieldTrip’s ‘ft_preproc_bandpassfilter’ with a two-pass FIR filter), followed by root mean square (RMS) calculation and smoothing with a 20-ms kernel (using MATLAB’s ‘smoothdata’ function). Sleep ripple events were identified when the smoothed RMS signal exceeded a threshold set at the mean plus three standard deviations (s.d.) of the RMS across all N2 and N3 data points. Events shorter than 38 ms (i.e., ≤ 3 cycles at 80 Hz) or longer than 300 ms were excluded. Time points exceeding an upper threshold (mean RMS + 9 s.d.) were also removed. The upward and downward crossings of the detection threshold determined the onset and offset of ripples. The ripple centers were defined as the maximal trough in the raw iEEG signal. Wake ripple events were detected using the same algorithm applied to wake time intervals. Surrogate ripple events were randomly sampled time points within the same vigilance state in a 5-minute time window around actual ripple events that contained no real ripple events or artifacts.

#### Ripple-triggered firing rates

Neuronal spiking activity was time-locked to the ripple center time, and instantaneous firing rates were calculated by binning neuronal spiking activity into 1 ms time bins, convolved with a 100 ms Gaussian kernel (smoothdata). The relative change in activity around ripples was obtained by averaging the ripple-triggered firing rate across all ripple events and subtracting the mean firing rate from a 1-second baseline window preceding the ripple (−3 to −2 seconds). Ripple-triggered firing rates were computed separately for ripples detected during wakefulness and sleep.

The population activity during ripples was obtained by averaging the ripple-triggered activity across all neurons. To compare the ripple-triggered activation during sleep versus wake ripple events, a paired-sample cluster permutation test comparing the activity within each neuron during wake and sleep ripples was used (*56*) (Edden M. Gerber 2025, permutest, MATLAB Central File Exchange, 10,000 permutations). Similarly, the time window of significant change to baseline firing during sleep and wake ripples was calculated by a paired-sample cluster permutation test against the baseline.

To ensure selective neurons reliably respond to their preferred stimulus and are tracked throughout the experiment, we performed two additional functional localizer sessions (‘screenings’), both before evening learning and after morning retrieval. During these localizers, all 12 stimuli were presented 10 times each for 1 s in pseudo-random order at screen center and participants answered a simple perceptual question about each image (e.g., “Is there a face present?”) to ensure attention. Neurons were classified as selective if they exhibited a significant firing rate increase to exactly one of the 12 images used in the learning task across both combined localizer sessions (evening & morning). Significance was defined relative to a 500 ms pre-stimulus baseline using an established bin-wise rank-sum test (*50*, *57*). Specifically, spiking activity during the 1-second image presentation was analyzed in 19 overlapping 100-ms bins (50-ms step width) and compared to the baseline spike distribution across all trials and stimuli (500-ms time window before stimulus onset) using a Wilcoxon rank-sum test. The resulting p-values were corrected for multiple testing across response bins using the Simes procedure (*57*, *58*). Only bins with firing rates above baseline were retained, and neurons were classified as selective if the resulting *P*_corrected_ < 0.05 for exactly one stimulus. Selective neurons were additionally required to be active in >50% of trials and show an average firing rate of ≥1 Hz during the response window. In total, 91 selective neurons were identified. These selective neurons provide a one-to-one mapping of neuronal activity and behavioral performance, offering the unique opportunity to probe the mechanisms of memory reactivation during human sleep by linking individual neurons to stimuli that are either remembered or forgotten. Images recalled correctly at both evening and morning retrieval were classified as successfully remembered, and all others were considered forgotten. Overall, 58 selective neurons responded to remembered items, and 33 to forgotten items.

To directly compare the ripple-triggered activity across neurons selective to remembered and forgotten items (Fig. 3), we normalized (z-scored) each neuron’s ripple-triggered firing rate using the mean and standard deviation of the 1-s baseline interval (−3 to −2 s prior to the ripple). This normalization controls for potential differences in baseline firing rate across neurons, ensuring that group-level comparisons reflect ripple-related modulation rather than intrinsic firing variability. Normalized ripple-triggered firing rates were then compared between neurons selective for remembered and forgotten items using a two-sample cluster permutation test (*56*) (permutest, 10,000 permutations).

#### Ripple-locked time-frequency representations

Time-frequency representations (TFRs) locked to ripples were computed for each macro electrode using the bipolar-referenced signals of the two innermost MTL contacts. First, the power spectra were calculated across the entire recording using Hanning tapers in 25 ms steps, with frequencies ranging from 1 to 200 Hz in 1 Hz increments. The taper window length was dynamically adjusted based on the frequency, spanning at least three cycles and at least 100 ms. Artifact-contaminated periods were padded by 200 ms, plus half the taper window length on both sides (with a minimum of 50 ms) and excluded from the analysis. The resulting power spectrum was normalized (z-scored) separately for wake and N2/3 sleep across all time bins per channel. For each macro contact, TFRs were separately segmented around detected ripple events during wakefulness and sleep and averaged using the trimmed mean to mitigate the influence of residual outliers by excluding the top and bottom 5%. The group ripple-triggered TFR was derived by first averaging the ripple-triggered TFRs across the innermost contact pairs in each session and then averaging across sessions. Ripple waveforms were extracted from the preprocessed iEEG data.

#### Identification and decoding of MTL bursts

Neuronal bursts are brief episodes in which a neuron fires multiple spikes in rapid succession, providing optimal conditions for synaptic plasticity and the coordinated transfer of information (*35*, *36*). Bursts were identified for each recorded neuron using non-parametric rank surprise statistics based on the ranks of inter-spike intervals across the entire spike train (*37*). Any spikes occurring within less than 200 ms of artifacts or interictal epileptiform discharges on the closest macro electrode were excluded from analysis. Compared to classical burst detection approaches such as the Poisson-surprise methods, the rank surprise is a robust approach which does not depend on prior assumptions about the distribution of inter-spike intervals. Potential burst events were required to contain a minimum of three spikes, with consecutive spikes occurring within less than 100 ms of each other. To capture only the most active bursts, a conservative burst detection threshold was applied (of RS^crit^ = 6) (*37*). Additionally, for each detected burst event, a surrogate burst event was identified by randomly selecting an artifact-free time point within ±30 seconds of the detected burst onset, that is at least 5 seconds away from the actual burst onset.

For each neuron, the relative burst rate around ripples was computed by averaging burst rates across all ripple events and subtracting the mean burst rate from a 10-second baseline window ending 3 seconds before each ripple. Burst rates were smoothed using a 100 ms Gaussian kernel (smoothdata). Ripple-aligned burst rates were calculated separately for wake and sleep. To compare burst activity across states, we performed a paired-sample cluster permutation test within each neuron, comparing burst rates during wake versus sleep ripples (*56*) (permutest, 10,000 permutations).

To test whether periods of MTL bursts can be identified in network-wide activity patterns, we first identified well-separated burst events (at least 2 seconds apart from other identified bursts) during both wakefulness and sleep (N2 & N3). To run the decoding, we required a minimum of 10 such events during wake and sleep periods. If more than 200 bursts were identified for a neuron, we randomly subsampled 200 events for computational efficiency. For each neuron, we extracted the burst-aligned iEEG signal from all macro contacts (80 to 96 contacts per participant). Line noise for each iEEG contact was removed using FieldTrip’s built-in function (ft_preproc_bandstopfilter, 4th order or reduced at 49–51, 99–101, 149–151, 199–201 Hz). The signal was high-pass filtered at 0.1 Hz (ft_preproc_highpassfilter, 4th order or reduced) and downsampled to 100 Hz. A 1-second time window around the burst onset was extracted, and the data were smoothed using the 100-ms moving mean (using MATLAB’s ‘movmean’ function). The resulting voltage traces were used to train linear multivariate pattern classifiers separately during sleep and wakefulness, using a linear discriminant analysis (LDA) implemented in the MVPA-Light toolbox (*59*). For decoding, data were first z-scored and averaged (2 trials), and a time-by-time decoding was performed across the entire 1-second time window using 5-fold cross-validation and 100 resample runs. Decoding performance during sleep and wakefulness was compared both against chance and between states using a paired-sample cluster permutation test (*56*) (permutest, 10,000 permutations).

## Supplementary figures

**Fig. S1.**
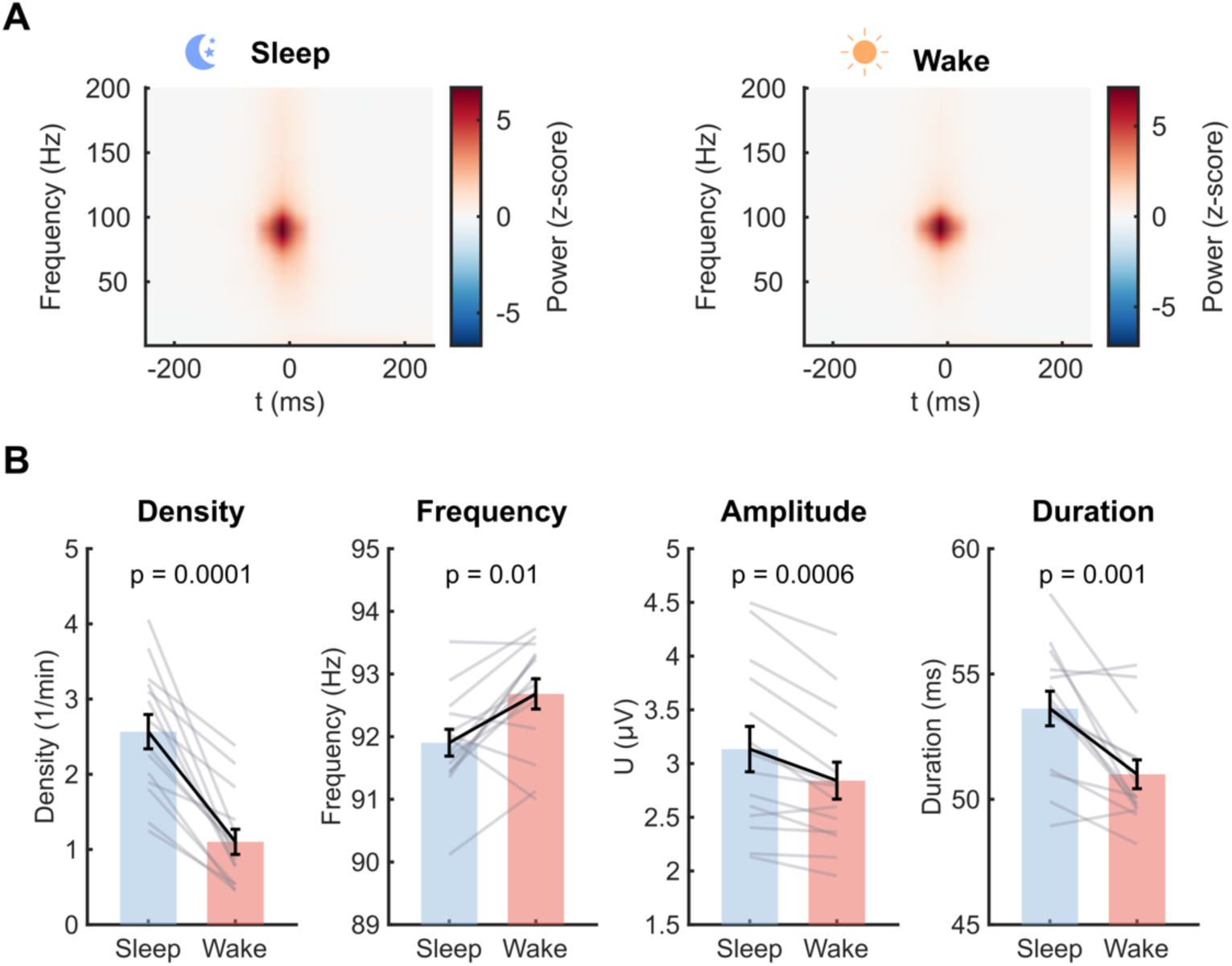
Ripple characteristics during wakefulness and sleep. **(A)** Time-frequency representations (TFRs, mean across 14 sessions) locked to the center of sleep (left) and wake (right) ripples. **(B)** Comparison of ripple density, central frequency, envelope amplitude, and duration (mean ± s.e.m.) during wakefulness and sleep across all 14 recording sessions. Ripples occurred significantly more often during sleep than wakefulness. Mean ripple frequency was higher in wakefulness, whereas sleep ripples exhibited greater envelope amplitude and longer duration (all tests based on N = 14 sessions and two-sided Wilcoxon signed-rank tests).

**Fig. S2.**
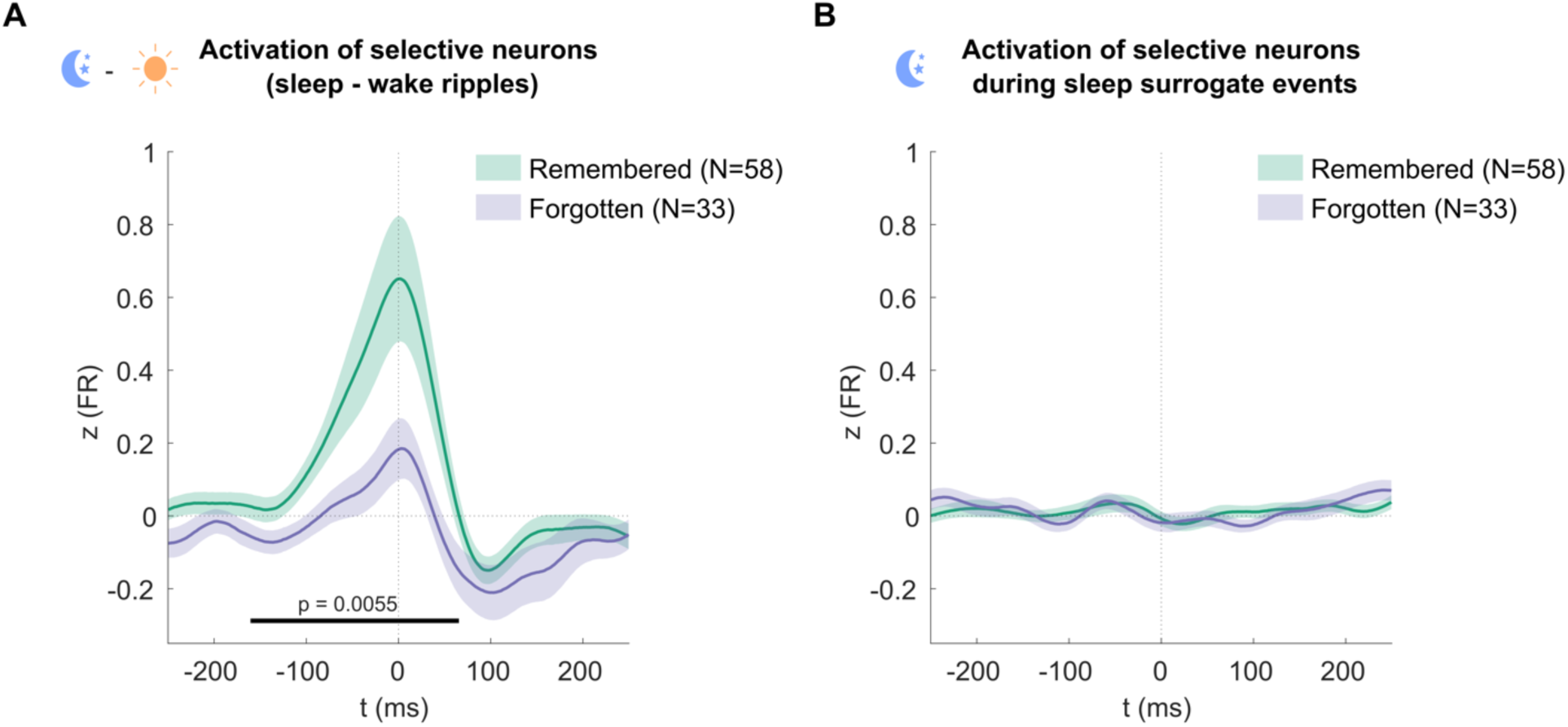
Sleep-specific ripple-triggered activation linked to memory performance. **(A)** Difference (mean ± s.e.m.) of wake and sleep ripple-triggered activity of neurons coding for remembered versus forgotten items (interaction; cluster permutation test; P = 0.0055; 10,000 permutations). **(B)** During ripple-free surrogate events, activity (mean ± s.e.m.) did not differ between neurons selective for remembered items and neurons selective for forgotten items (cluster permutation test; 10,000 permutations).

**Fig. S3.**
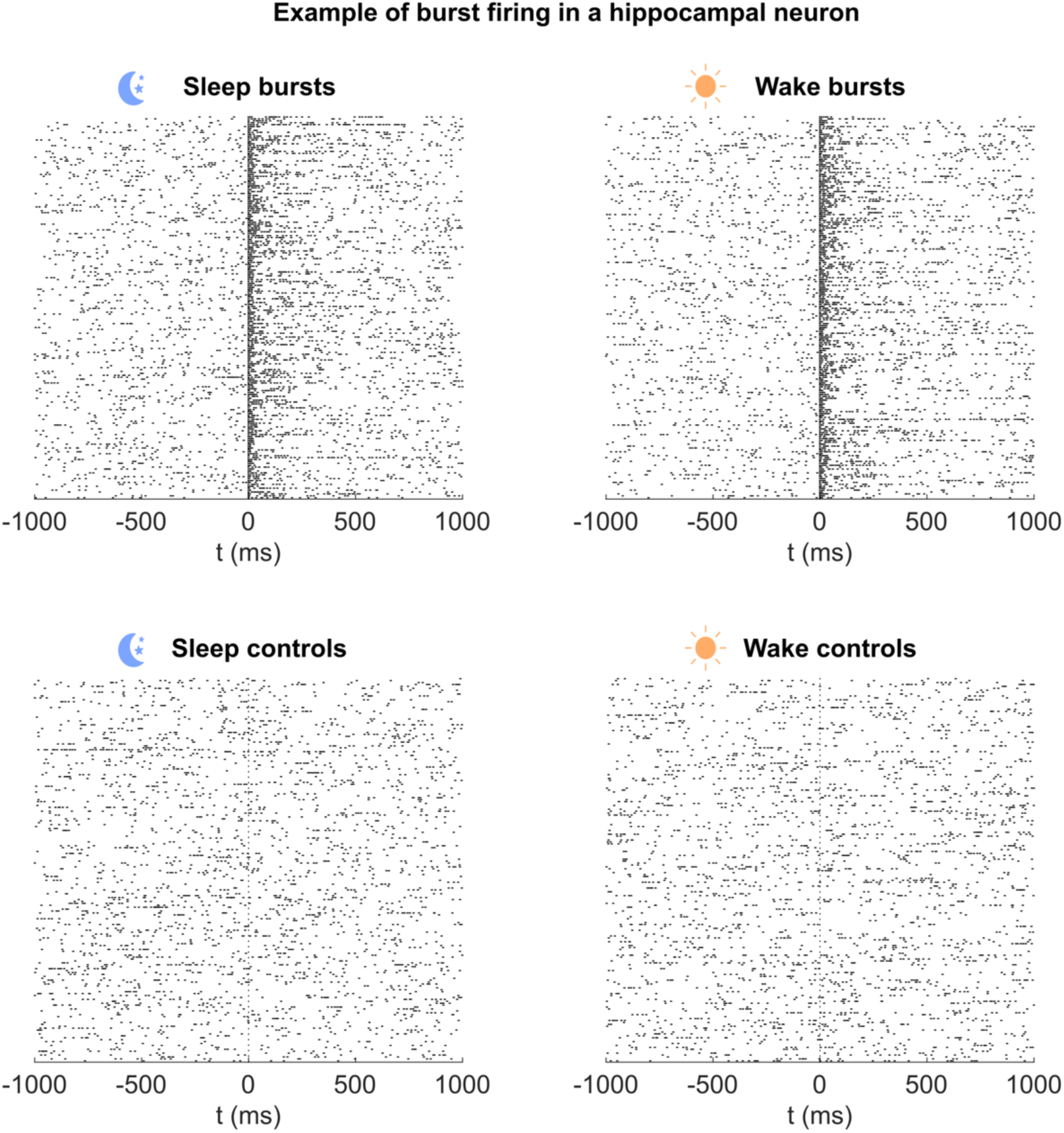
Examples of detected neuronal burst events. Top row: Raster plots aligned to burst onset for a hippocampal neuron, showing a subset of 200 bursts detected during sleep (left) and wakefulness (right). Bursts were detected independently of ripples, for each neuron. Bottom row: Raster plots aligned to surrogate non-burst events selected within the temporal proximity of detected burst events, shown separately for sleep (left) and wakefulness (right).

**Table S1.**
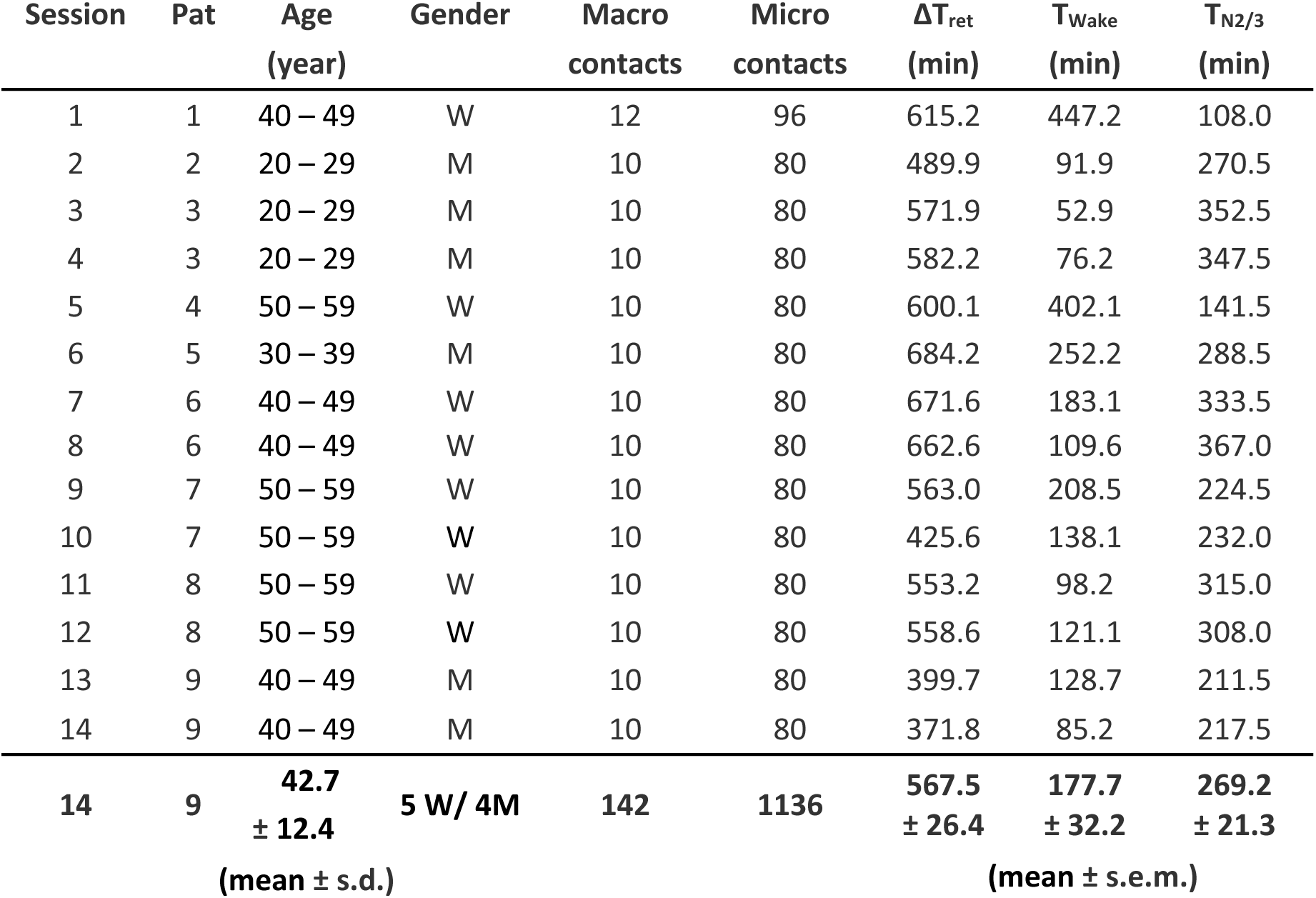
Overview of recording sessions and participant characteristics. All patients were bilaterally implanted in the medial temporal lobe (MTL) with Behnke-Fried electrodes for invasive pre-surgical epilepsy monitoring. Each MTL electrode contained eight clinical macro contacts and one microwire bundle comprising eight recording microwires. ΔT_ret_ denotes the total interval between evening and morning retrieval, T_wake_ the cumulative time spent awake between retrieval sessions, and T_N2/3_ the time spent in non-REM sleep stages 2 and 3. Age information is reported in ranges to protect patient confidentiality.

## References and Notes

1. J. G. Klinzing, N. Niethard, J. Born, Mechanisms of systems memory consolidation during sleep. Nat. Neurosci. 22, 1598–1610 (2019).

2. B. Rasch, J. Born, About Sleep’s Role in Memory. Physiol. Rev. 93, 681–766 (2013).

3. B. P. Staresina, Coupled sleep rhythms for memory consolidation. Trends Cogn. Sci. 28, 339–351 (2024).

4. G. Buzsáki, Hippocampal sharp wave-ripple: A cognitive biomarker for episodic memory and planning. Hippocampus 25, 1073–1188 (2015).

5. Z. S. Chen, M. A. Wilson, How our understanding of memory replay evolves. J. Neurophysiol. 129, 552–580 (2023).

6. K. Diba, G. Buzsáki, Forward and reverse hippocampal place-cell sequences during ripples. Nat. Neurosci. 10, 1241–1242 (2007).

7. M. A. Wilson, B. L. McNaughton, Reactivation of Hippocampal Ensemble Memories During Sleep. Science 265, 676–679 (1994).

8. G. Girardeau, K. Benchenane, S. I. Wiener, G. Buzsáki, M. B. Zugaro, Selective suppression of hippocampal ripples impairs spatial memory. Nat. Neurosci. 12, 1222–1223 (2009).

9. E. Tulving, Episodic Memory: From Mind to Brain. Annu. Rev. Psychol. 53, 1–25 (2002).

10. M. Schönauer, S. Alizadeh, H. Jamalabadi, A. Abraham, A. Pawlizki, S. Gais, Decoding material-specific memory reprocessing during sleep in humans. Nat. Commun. 8, 15404 (2017).

11. T. Schreiner, M. Petzka, T. Staudigl, B. P. Staresina, Respiration modulates sleep oscillations and memory reactivation in humans. Nat. Commun. 14, 8351 (2023).

12. T. Schreiner, M. Petzka, T. Staudigl, B. P. Staresina, Endogenous memory reactivation during sleep in humans is clocked by slow oscillation-spindle complexes. Nat. Commun. 12, 3112 (2021).

13. B. P. Staresina, A. Alink, N. Kriegeskorte, R. N. Henson, Awake reactivation predicts memory in humans. Proc. Natl. Acad. Sci. 110, 21159–21164 (2013).

14. A. Tambini, L. Davachi, Awake Reactivation of Prior Experiences Consolidates Memories and Biases Cognition. Trends Cogn. Sci. 23, 876–890 (2019).

15. A. Bragin, J. Engel Jr, C. L. Wilson, I. Fried, G. Buzsáki, High-frequency oscillations in human brain. Hippocampus 9, 137–142 (1999).

16. R. F. Helfrich, J. D. Lendner, B. A. Mander, H. Guillen, M. Paff, L. Mnatsakanyan, S. Vadera, M. P. Walker, J. J. Lin, R. T. Knight, Bidirectional prefrontal-hippocampal dynamics organize information transfer during sleep in humans. Nat. Commun. 10, 3572 (2019).

17. X. Jiang, J. Gonzalez-Martinez, E. Halgren, Posterior Hippocampal Spindle Ripples Co-occur with Neocortical Theta Bursts and Downstates-Upstates, and Phase-Lock with Parietal Spindles during NREM Sleep in Humans. J. Neurosci. 39, 8949–8968 (2019).

18. B. P. Staresina, J. Niediek, V. Borger, R. Surges, F. Mormann, How coupled slow oscillations, spindles and ripples coordinate neuronal processing and communication during human sleep. Nat. Neurosci., 1–9 (2023).

19. B. P. Staresina, T. O. Bergmann, M. Bonnefond, R. van der Meij, O. Jensen, L. Deuker, C. E. Elger, N. Axmacher, J. Fell, Hierarchical nesting of slow oscillations, spindles and ripples in the human hippocampus during sleep. Nat. Neurosci. 18, 1679–1686 (2015).

20. S. P. Jadhav, C. Kemere, P. W. German, L. M. Frank, Awake Hippocampal Sharp-Wave Ripples Support Spatial Memory. Science 336, 1454–1458 (2012).

21. J. O’Neill, B. Pleydell-Bouverie, D. Dupret, J. Csicsvari, Play it again: reactivation of waking experience and memory. Trends Neurosci. 33, 220–229 (2010).

22. J. O’Neill, T. Senior, J. Csicsvari, Place-Selective Firing of CA1 Pyramidal Cells during Sharp Wave/Ripple Network Patterns in Exploratory Behavior. Neuron 49, 143–155 (2006).

23. Y. Y. Chen, L. Aponik-Gremillion, E. Bartoli, D. Yoshor, S. A. Sheth, B. L. Foster, Stability of ripple events during task engagement in human hippocampus. Cell Rep. 35, 109304 (2021).

24. R. Q. Quiroga, L. Reddy, G. Kreiman, C. Koch, I. Fried, Invariant visual representation by single neurons in the human brain. Nature 435, 1102–1107 (2005).

25. M. S. Kehl, S. Mackay, K. Ohla, M. Schneider, V. Borger, R. Surges, M. Spehr, F. Mormann, Single-neuron representations of odours in the human brain. Nature 634, 626–634 (2024).

26. R. Quian Quiroga, 20 years of concept cells: From invariant responses to a unique coding of human memory. Neuron, S0896-6273(26)00051–6 (2026).

27. R. Q. Quiroga, A. Kraskov, C. Koch, I. Fried, Explicit encoding of multimodal percepts by single neurons in the human brain. Curr. Biol. CB 19, 1308–1313 (2009).

28. M. Bausch, J. Niediek, T. P. Reber, S. Mackay, J. Boström, C. E. Elger, F. Mormann, Distinct neuronal populations in the human brain combine content and context. Nature 650, 690–700 (2026).

29. R. Quian Quiroga, Concept cells: the building blocks of declarative memory functions. Nat Rev Neurosci 13, 587–597 (2012).

30. S. Mackay, T. P. Reber, M. Bausch, J. Boström, C. E. Elger, F. Mormann, Concept and location neurons in the human brain provide the “what” and “where” in memory formation. Nat. Commun. 15, 7926 (2024).

31. T. V. P. Bliss, T. Lømo, Long-lasting potentiation of synaptic transmission in the dentate area of the anaesthetized rabbit following stimulation of the perforant path. J. Physiol. 232, 331–356 (1973).

32. U. Frey, R. G. M. Morris, Synaptic tagging and long-term potentiation. Nature 385, 533–536 (1997).

33. Y. Dudai, A. Karni, J. Born, The Consolidation and Transformation of Memory. Neuron 88, 20–32 (2015).

34. P. W. Frankland, B. Bontempi, The organization of recent and remote memories. Nat. Rev. Neurosci. 6, 119–130 (2005).

35. J. E. Lisman, Bursts as a unit of neural information: making unreliable synapses reliable. Trends Neurosci. 20, 38–43 (1997).

36. A. Payeur, J. Guerguiev, F. Zenke, B. A. Richards, R. Naud, Burst-dependent synaptic plasticity can coordinate learning in hierarchical circuits. Nat. Neurosci. 24, 1010–1019 (2021).

37. B. Gourévitch, J. J. Eggermont, A nonparametric approach for detection of bursts in spike trains. J. Neurosci. Methods 160, 349–358 (2007).

38. J. Niediek, T. P. Reber, M. Bausch, F. Schwimmbeck, H. Gast, V. A. Coenen, J. Boström, C. E. Elger, F. Mormann, Episodic memory consolidation by reactivation of human concept neurons during sleep reflects contents, not sequence of events. bioRxiv [Preprint] (2026). 10.64898/2026.01.10.698827.

39. B. Rasch, C. Büchel, S. Gais, J. Born, Odor Cues During Slow-Wave Sleep Prompt Declarative Memory Consolidation. Science 315, 1426–1429 (2007).

40. N. W. Schuck, Y. Niv, Sequential replay of nonspatial task states in the human hippocampus. Science 364 (2019).

41. G. Buzsáki, Z. Horváth, R. Urioste, J. Hetke, K. Wise, High-frequency network oscillation in the hippocampus. Science 256, 1025–1027 (1992).

42. J. G. Jenkins, K. M. Dallenbach, Obliviscence during Sleep and Waking. Am. J. Psychol. 35, 605–612 (1924).

43. G. Tononi, C. Cirelli, Sleep and the Price of Plasticity: From Synaptic and Cellular Homeostasis to Memory Consolidation and Integration. Neuron 81, 12–34 (2014).

44. S. Cassenaer, G. Laurent, Conditional modulation of spike-timing-dependent plasticity for olfactory learning. Nature 482, 47–52 (2012).

45. L. Nadel, M. Moscovitch, Memory consolidation, retrograde amnesia and the hippocampal complex. Curr. Opin. Neurobiol. 7, 217–227 (1997).

46. J. Born, I. Wilhelm, System consolidation of memory during sleep. Psychol. Res. 76, 192–203 (2012).

47. A. Fernández-Ruiz, A. Oliva, E. Fermino de Oliveira, F. Rocha-Almeida, D. Tingley, G. Buzsáki, Long-duration hippocampal sharp wave ripples improve memory. Science 364, 1082–1086 (2019).

48. J. Niediek, J. Boström, C. E. Elger, F. Mormann, Reliable Analysis of Single-Unit Recordings from the Human Brain under Noisy Conditions: Tracking Neurons over Hours. PLoS ONE 11 (2016).

49. M. Bausch, J. Niediek, T. P. Reber, S. Mackay, J. Boström, C. E. Elger, F. Mormann, Concept neurons in the human medial temporal lobe flexibly represent abstract relations between concepts. Nat. Commun. 12, 6164 (2021).

50. T. P. Reber, S. Mackay, M. Bausch, M. S. Kehl, V. Borger, R. Surges, F. Mormann, Single-neuron mechanisms of neural adaptation in the human temporal lobe. Nat. Commun. 14, 2496 (2023).

51. R. Oostenveld, P. Fries, E. Maris, J.-M. Schoffelen, FieldTrip: Open Source Software for Advanced Analysis of MEG, EEG, and Invasive Electrophysiological Data. Comput. Intell. Neurosci. 2011, 156869 (2011).

52. R. Janca, P. Jezdik, R. Cmejla, M. Tomasek, G. A. Worrell, M. Stead, J. Wagenaar, J. G. R. Jefferys, P. Krsek, V. Komarek, P. Jiruska, P. Marusic, Detection of Interictal Epileptiform Discharges Using Signal Envelope Distribution Modelling: Application to Epileptic and Non-Epileptic Intracranial Recordings. Brain Topogr. 28, 172–183 (2015).

53. R. Vallat, M. P. Walker, An open-source, high-performance tool for automated sleep staging. eLife 10, e70092 (2021).

54. A. Guillot, V. Thorey, RobustSleepNet: Transfer Learning for Automated Sleep Staging at Scale. IEEE Trans. Neural Syst. Rehabil. Eng. 29, 1441–1451 (2021).

55. H.-V. Ngo, J. Fell, B. Staresina, Sleep spindles mediate hippocampal-neocortical coupling during long-duration ripples. eLife 9, e57011 (2020).

56. E. Maris, R. Oostenveld, Nonparametric statistical testing of EEG- and MEG-data. J. Neurosci. Methods 164, 177–190 (2007).

57. F. Mormann, S. Kornblith, R. Q. Quiroga, A. Kraskov, M. Cerf, I. Fried, C. Koch, Latency and Selectivity of Single Neurons Indicate Hierarchical Processing in the Human Medial Temporal Lobe. J. Neurosci. 28, 8865–8872 (2008).

58. R. J. Simes, An improved Bonferroni procedure for multiple tests of significance. Biometrika 73, 751–754 (1986).

59. M. S. Treder, MVPA-Light: A Classification and Regression Toolbox for Multi-Dimensional Data. Front. Neurosci. 14 (2020).

